# TMPRSS2 inhibitor discovery facilitated through an *in silico* and biochemical screening platform

**DOI:** 10.1101/2021.03.22.436465

**Authors:** Amanda L. Peiffer, Julie M. Garlick, Yujin Wu, Matthew B. Soellner, Charles L. Brooks, Anna K. Mapp

## Abstract

The COVID-19 pandemic has highlighted the need for new antiviral targets, as many of the currently approved drugs have proven ineffective against mitigating SARS-CoV-2 infections. The host transmembrane serine protease TMPRSS2 is a highly promising antiviral target, as it plays a direct role in priming the spike protein before viral entry occurs. Further, unlike other targets such as ACE2, TMPRSS2 has no known biological role. Here we utilize virtual screening to curate large libraries into a focused collection of potential inhibitors. Optimization of a recombinant expression and purification protocol for the TMPRSS2 peptidase domain facilitates subsequent biochemical screening and characterization of selected compounds from the curated collection in a kinetic assay. In doing so, we demonstrate that serine protease inhibitors camostat, nafamostat, and gabexate inhibit through a covalent mechanism. We further identify new non-covalent compounds as TMPRSS2 protease inhibitors, demonstrating the utility of a combined virtual and experimental screening campaign in rapid drug discovery efforts.

## Introduction

The emergence of COVID-19 in late 2019 and the rapid transmission of the disease around the globe has prompted an urgent need for effective treatments.^1^ As with many coronoviruses, infection with SARS-CoV-2 requires host cell cooperation; the spike (S) protein protruding outside the viral coat requires priming by TMPRSS2, a human transmembrane serine protease, for viral entry via the receptor ACE2 (Figure 1A).^2–5^ While many have focused on blocking the interactions between ACE2 and the S protein, ACE2 also plays an important role in healthy cell function by counterbalancing ACE to lower and maintain healthy blood pressure.^6^ Alternatively, there is little known about the biological function of TMPRSS2, with data suggesting it is likely functionally redundant.^7,8^ Along with SARS-CoV-2, TMPRSS2 has been implicated in priming other pathogenic coronaviruses such as SARS-CoV and MERS, as well as influenza.^9–12^ TMPRSS2^−/−^ knockout mice have little phenotypic differences compared to wild-type animals, yet conferred resistance to viral infections, suggesting that the protein is not essential.^13^ Inhibiting transcription of TMPRSS2 via BET inhibitors leads to decreased infectivity of SARS-CoV-2 in human lung cells, further suggesting the viability of TMPRSS2 inhibition as an antiviral strategy.^14^ Additionally, as a human protein target rather than a viral protein target, TMPRSS2-targeting therapeutics should be less susceptible to drug-resistance due to viral mutation. Thus, TMPRSS2 is a desirable drug target for treating SARS-CoV-2 and future coronavirus infections.

**Figure 1.**
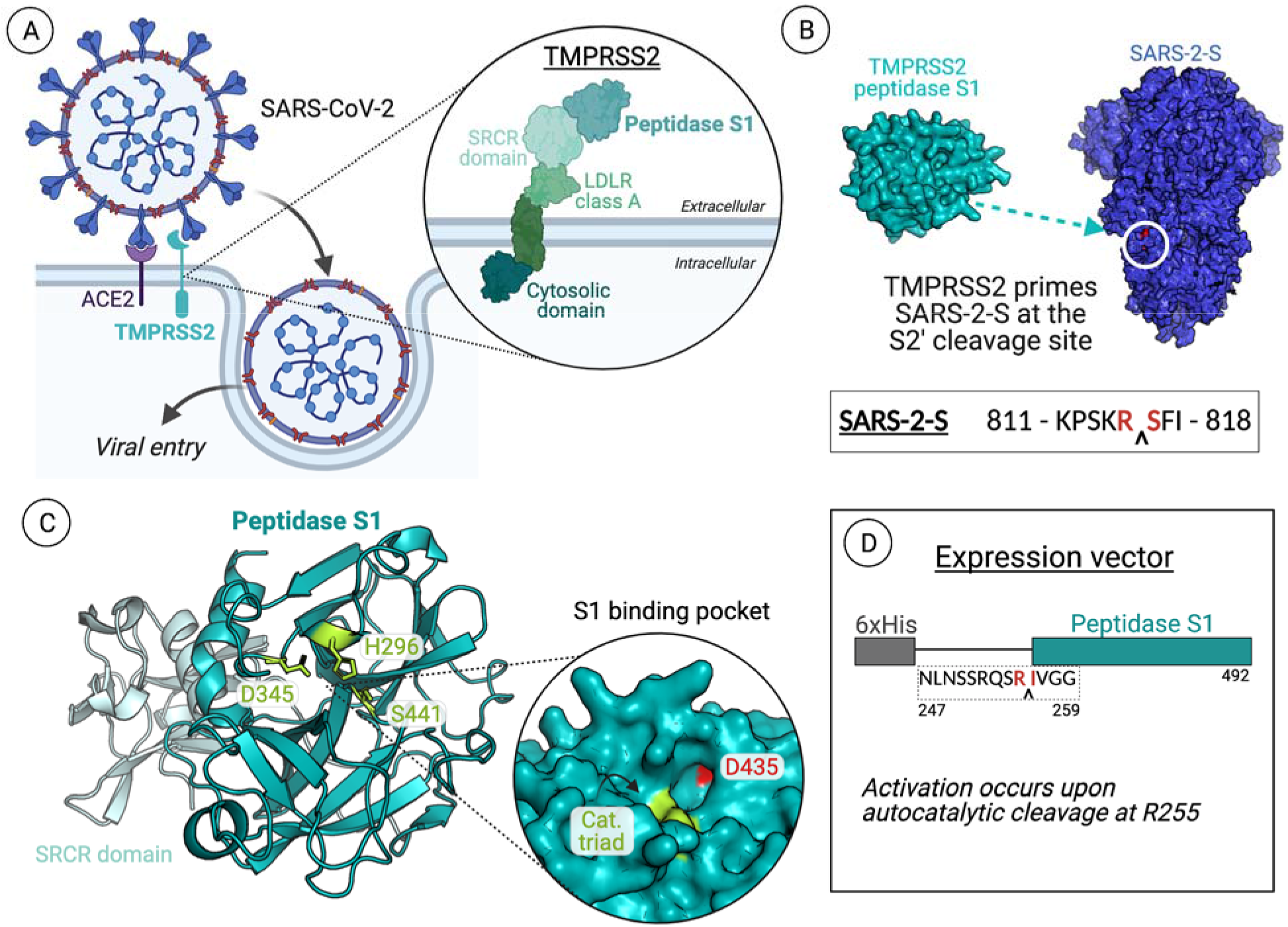
The role of TMPRSS2 in SARS-CoV-2 infection. (A) TMPRSS2 primes the viral S protein (SARS-2-S), which promotes membrane fusion and ultimately viral entry. TMPRSS2, part of the type II transmembrane serine protease family and hepsin/TMPRSS subfamily, is anchored at the cell membrane.^4^ The protein is mostly extracellular, with a small intracellular cytosolic domain. The extracellular portion of the protein is composed of a LDLR class A domain, an SRCR domain, and finally the peptidase S1 domain required for catalytic activity.^23^ (B) The peptidase S1 domain of TMPRSS2 cleaves SARS-2-S at the S2’ cleavage site.^24,25^ (C) Homology model of TMPRSS2 SRCR and peptidase S1 domains, generated using SWISS-Model. Peptidase S1 domains are chymotrypsin-like and contain the catalytic triad histidine, aspartic acid, and serine (active site residues for TMPRSS2 are H296, D345, S441).^23^ (D) The DNA vector constructed for recombinant expression and purification for the protease domain of TMPRSS2. Figure created using BioRender.com.

To date, there are few reported TMPRSS2 inhibitors. Camostat, a compound initially discovered as a Matriptase 2 inhibitor, also inhibits TMPRSS2.^4,15^ Nafamostat and gabexate have also been reported to inhibit TMPRSS2. However, both camostat and nafamostat inhibit a wide range of serine proteases and are rapidly metabolized in mammals to structures with poorly defined activity. ^16–18^ It has been reported that each of these compounds all form a covalent bond with the active site serine of serine proteases via the central ester, also a site of metabolic breakdown (Figure 2A).^19,20^ Additionally, molecular modeling studies on TMPRSS2 supports covalent bond formation with both camostat and nafamostat, as well as the camostat metabolite FOY 251.^21^ Thus, TMPRSS2 inhibitors with less reactive architectures are highly desirable.

**Figure 2.**
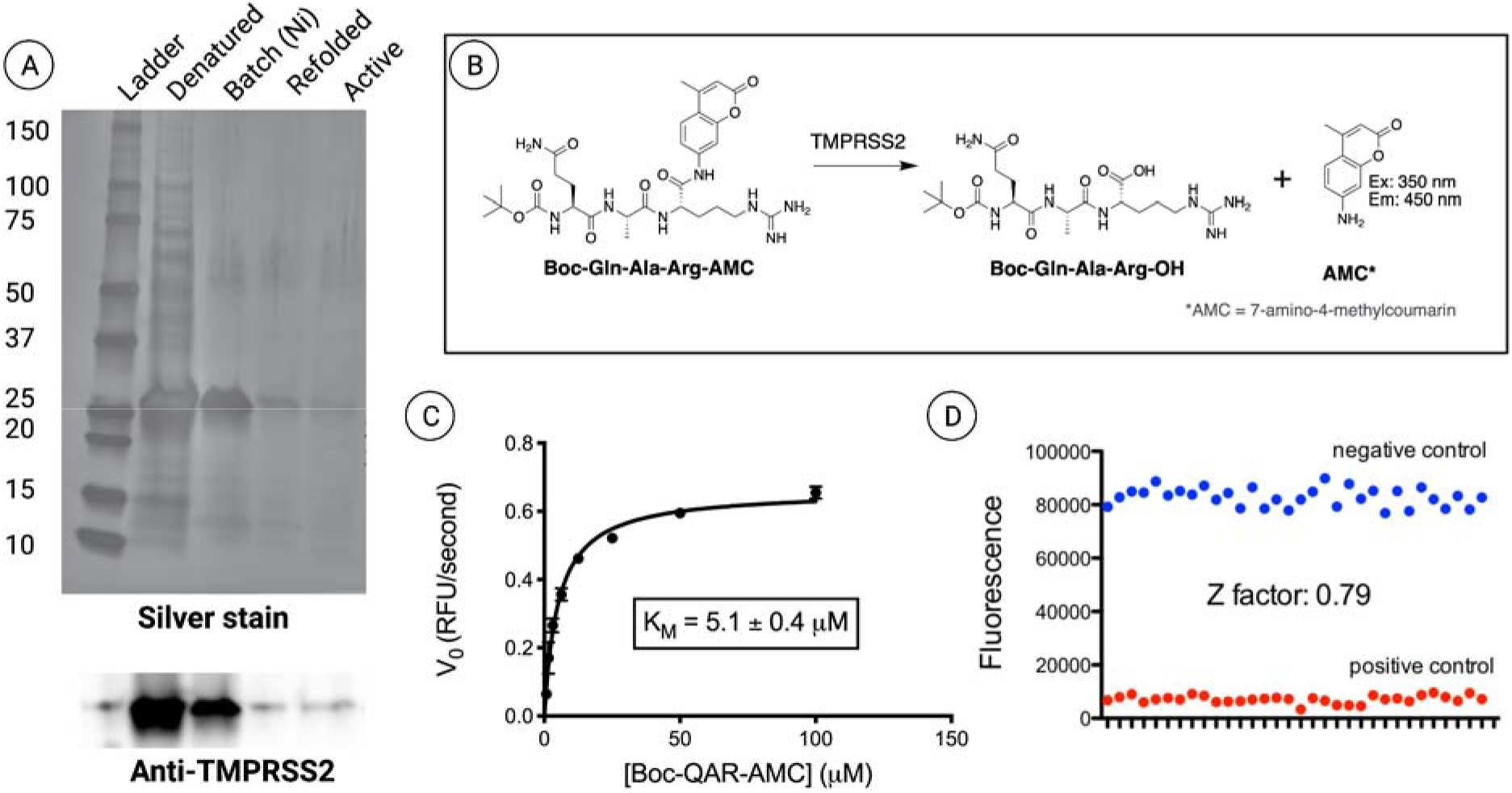
(A) TMPRSS2 protein is expressed and purified by expression in inclusion bodies (labeled denatured), purified by batch binding to Ni-NTA resin, refolded, and dialyzed into assay buffer. Silver staining shows purity and protein levels at each step (TMPRSS2, ~26kDa). Western blot with TMPRSS2 antibody raised against the protease domain confirms the identity of the 26 kDa protein observed with silver staining as TMPRSS2. (B) Schematic of the biochemical assay used to monitor TMPRSS2 activity. (C) K_M_ of Boc-Glu-Ala-Arg-AMC is determined by plotting initial velocity, calculated as the slope of the kinetic trace at less than 10 percent substrate cleavage, vs substrate concentration. Data was fit to the Michaelis-Menten equation using GraphPad Prism. (D) TMPRSS2 activity assay was adapted to 384 well format with the Z factor indicating an excellent assay. Negative controls correspond to protein + substrate, while positive controls correspond to substrate only.

As a strategy for rapid TMPRSS2 inhibitor discovery, we developed a combined *in silico* and biochemical workflow. Strategic development of a TMPRSS2 expression protocol, utilizing an autocatalysis-based affinity tag removal, facilitated purification for biochemical assay development. This allowed existing TMPRSS2 inhibitors to be profiled and characterized as covalent. Our protocol for virtual screening against a TMPRSS2 homology model (Figure 1C) comprises a molecular dynamics/simulated annealing-based docking that employs flexible receptor side chains to capture subtle changes, both conformational and energetic, for improved compound scoring. Our approach curated a subset of promising TMPRSS2 ligands, which upon subsequent biochemical testing were identified as active inhibitors. In doing so, we identify new non-covalent hit compounds that can be both repurposed for SARS-CoV-2 infections as well as derivatized to yield improved TMPRSS2 inhibitors.

## Results

### Active TMPRSS2 Peptidase S1 can be expressed recombinantly in *E. coli*

Like other serine proteases, TMPRSS2 is natively expressed as a zymogen; activation occurs via autocatalysis of the peptide bond between Arg-255 and Ile-256, leaving the *N*-terminal portion of Ile-256 to undergo a conformational change to stabilize the active state.^26–28^ Careful consideration was taken when designing the gene fragment for recombinant expression and purification of the TMPRSS2 peptidase domain (residues 256–492). *C*-terminal affinity tags appear to disrupt catalytic activity; thus, an *N*-terminal affinity tag is required but must be removed, leaving residue Ile-256 with a free *N*-terminus. Previous approaches to expression and purification in bacteria have utilized an orthogonal protease to allow for cleavage directly *N*-terminal to Ile-256, such as the TAGzyme system.^22^ We hypothesized that utilizing a construct with an extended *N*-terminal portion would be enough to promote enzyme autocatalysis, removing any *N*-terminal affinity tag as well as yielding the necessary free isoleucine without the need for multiple rounds of purification and introduction of an orthogonal cleavage site. Thus, a gene fragment for the catalytic domain of TMPRSS2 was constructed (see Methods), called TMPRSS2(247–492), with an *N*-terminal 6xHis tag for purification (Figure 1D).

Overexpression of the TMPRSS2(247–492) construct in *E. coli* led to protein aggregation in insoluble inclusion bodies, which enabled simple separation from the remaining cell lysate. Solubilizing the inclusion bodies required denaturing the aggregates utilizing 8 M urea. The N-terminal 6xHis tag was used to remove the denatured, unfolded protein from remaining impurities by batch binding with Ni NTA resin. Purified, denatured TMPRSS2 was then subjected to refolding by rapid dilution in 1:100 refolding buffer. Development of a modified refolding procedure using a syringe pump for slow and controlled dilution proved to be instrumental in producing active protein, likely due to providing a more optimal environment for the three internal disulfide bonds to correctly form. Concentration (10-fold) and subsequent dialysis into a 50 mM Tris 500 mM NaCl 0.01% NP-40 pH 8 buffer led to activation of the enzymatic activity demonstrated by the autocatalytic cleavage of the 6xHis tag (Figure S1).

### Activity of TMPRSS2 peptidase domain

The classic trypsin substrate Boc-QAR-AMC has been reported as a TMPRSS2 substrate, which we used to confirm activity of the purified protein (Figure 2B).^16,29^ To determine approximate concentration of active protein, we measured TMPRSS2 activity while titrating in the covalent protease inhibitor FPR-chloromethylketone (also known as PPACK).^30–32^ This inhibitor reacts with the histidine in the catalytic triad of serine proteases to irreversibly alkylate and inactivate the enzyme in a 1:1 complex (Figure S2). We determined the K_M_ to be 5.1 ± 0.4 μM with TMPRSS2 (Figure 2C), which is comparable to the K_M_ for trypsin with this substrate, 7.9 ± 0.8 μM (Figure S3).

Experimental conditions for high throughput screening (HTS) in 384-well plates were optimized using 2.5 μM substrate and 0.5 nM TMPRSS2. The concentration of substrate in this assay was set below the K_M_ to enable identification of competitive inhibitors. After incubation at room temperature for 30 min, endpoint fluorescence was determined (ex: 355 nm, em: 450 nm). At 30 min, less than 20% of the substrate was cleaved as determined by comparing fluorescence to that of 2.5 μM free AMC (i.e. 100% substrate conversion), enabling inhibition to be monitored while activity is still within the linear range. The S:B and Z factor were calculated to be 10.6 and 0.79, respectively, indicating an excellent HTS assay (Figure 2D). This biochemical assay was then utilized to test the apparent IC_50_ of known inhibitors camostat, nafamostat, and gabexate, where the compounds were incubated with TMPRSS2 for 30 min before substrate was added (Figure S4). IC_50_ values for these compounds agree with previously reported values obtained using similar biochemical approaches. Notably, the IC_50_ values for these compounds are very low, approaching the lower limit of detection based on the protein concentration in the assay.

### Analysis of covalent TMPRSS2 inhibitors

Although not previously shown experimentally, others have suggested that the inhibitors camostat and nafamostat covalently modify TMPRSS2 with a similar mechanism to other proteases.^21^ This involves initial binding, acylation, and ultimately hydrolysis. While it would be ideal to validate formation of a covalent adduct with TMPRSS2 using mass spectrometry, the purification of the protease domain yields low concentrations of active protein that is very sensitive to buffer conditions, making mass spectrometric analysis intractable. However, we demonstrate covalent adduct formation via mass spectrometry using trypsin (Figure 3B). With equimolar protein and compound, an increase in mass of 161 Da is observed for camostat, nafamostat, and FOY 251 (Figure S5). Gabexate requires 10X compound compared to protein for adduct formation to be observed, which corresponds to a mass increase of 155 Da (Figure S6). Additionally, a time-dependent decrease in apparent IC_50_ is observed with camostat in biochemical activity assays for both trypsin and TMPRSS2 (Figure 3C). This shift in IC_50_ vs. time fits to an equation for one phase decay, with the IC_50_ values approaching the limiting concentration of the protein used in the assay, suggesting that covalent bond formation is occurring at the active site (Figure 5A). Thus, while these compounds exhibit low nM IC_50_ values *in vitro*, this is likely due, at least in part, to the covalent mechanism of action.

**Figure 3.**
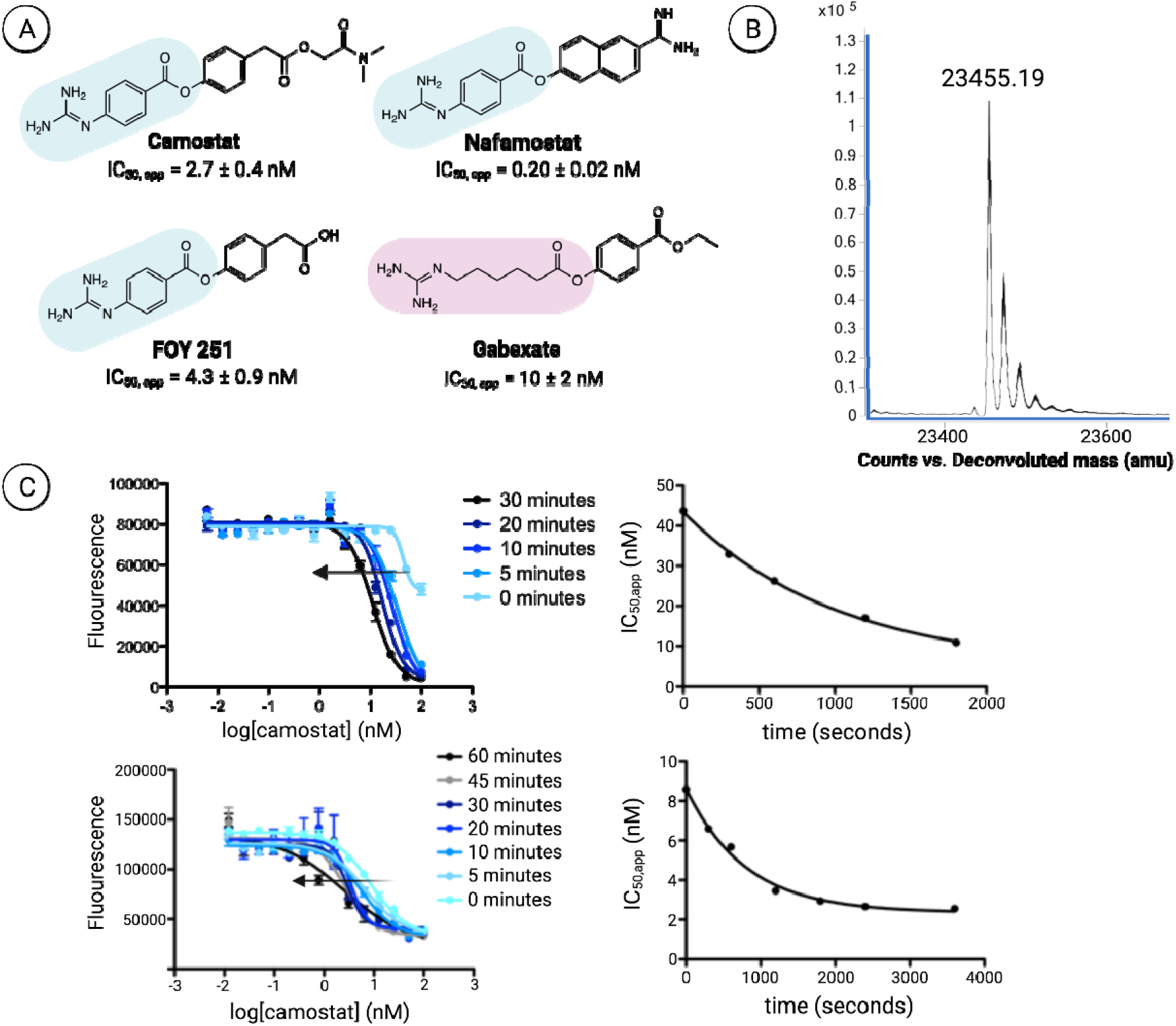
(A) The molecules that have commonly been cited as TMPRSS2 inhibitors are camostat, FOY 251, nafamostat, and gabexate. All contain a reactive ester, which can form a covalent bond with the catalytic serine. Apparent IC_50_ values for the four inhibitors against TMPRSS2 after 30 minutes incubation time are shown.(B) Deconvoluted mass spec data for trypsin incubated with camostat for 30 minutes shows formation of the covalent adduct. (C) Decreasing IC_50_ values for camostat with increased incubation time for trypsin (top) and TMPRSS2 (bottom) is consistent with covalent inhibition. Figure created using BioRender.com.

It is important to note that in a cellular context, camostat has a very short half-life of <1 min.^33^ Rapid hydrolysis to (4◻(4◻guanidinobenzoyl◻oxy)phenylacetic acid), also known as the protease inhibitor FOY 251 (Figure 3A), occurs both *in vitro* in serum as well as *in vivo*.^34–36^ We observe similar potencies of FOY 251 and camostat in our biochemical assay, with apparent IC_50_ = 4.3 ± 0.9 nM and 2.7 ± 0.4 nM, respectively (Figure S4). The half◻life of FOY 251 is longer than camostat, though it is metabolized to 4-guanidinobenzoic acid, which shows minimal TMPRSS2 inhibition.^34^ Thus, it would be advantageous to identify novel inhibitors that have alternative molecular scaffolds.

### Construction and refinement of TMPRSS2 homology model

Virtual screening methods have greatly improved over the past two decades, leading many drug discovery campaigns by filtering out thousands/millions of molecules before testing them *in vitro*. However, such studies require a structural model in which to dock compounds into the active site. Because no crystallographic or NMR-based models exist for TMPRSS2, we developed a homology model for the active soluble domain starting from prediction using the SWISS-MODEL web-interface (Figure 1C).^37–41^ This structure was built based on sequence homology of hepsin (PDB 5CE1). The structure showed high homology with the TMPRSS2 peptidase domain (34% sequence similarity with 70% sequence coverage) and also contained the bound ligand 2-[6-(1-hydroxycyclohexyl)pyridin-2-yl]-1H-indole-5-carboximidamide, which served as one of the templates for pharmacophore-based docking of putative ligands as described below. The SWISS-MODEL structure of TMPRSS2 was further “conditioned” through the application of molecular dynamics in an implicit solvent (GBMV) model to facilitate better packing and configurational relaxation.^42–44^

### Virtual screening yields preliminary hits for *in vitro* assays

Extensive virtual screening was performed to obtain putative hits for follow-up testing via *in vitro* inhibition assays (Figure 4). A total of 134,109 molecules were collected from multiple databases, which were subjected to a hierarchical refinement of docking poses. In the first stage, rigid receptor docking was performed exploring two means of initially positioning the small molecules. One utilized pharmacophores based on ligands in other bound serine proteases (see Methods), and the other initiated from a random generation of molecular conformations and random positioning inside the pocket (Figure 4A). The second relied upon a novel 3D pharmacophore *fastdock* framework, which operates by superposing pharmacophores onto compounds bound in experimentally solved structures.

**Figure 4.**
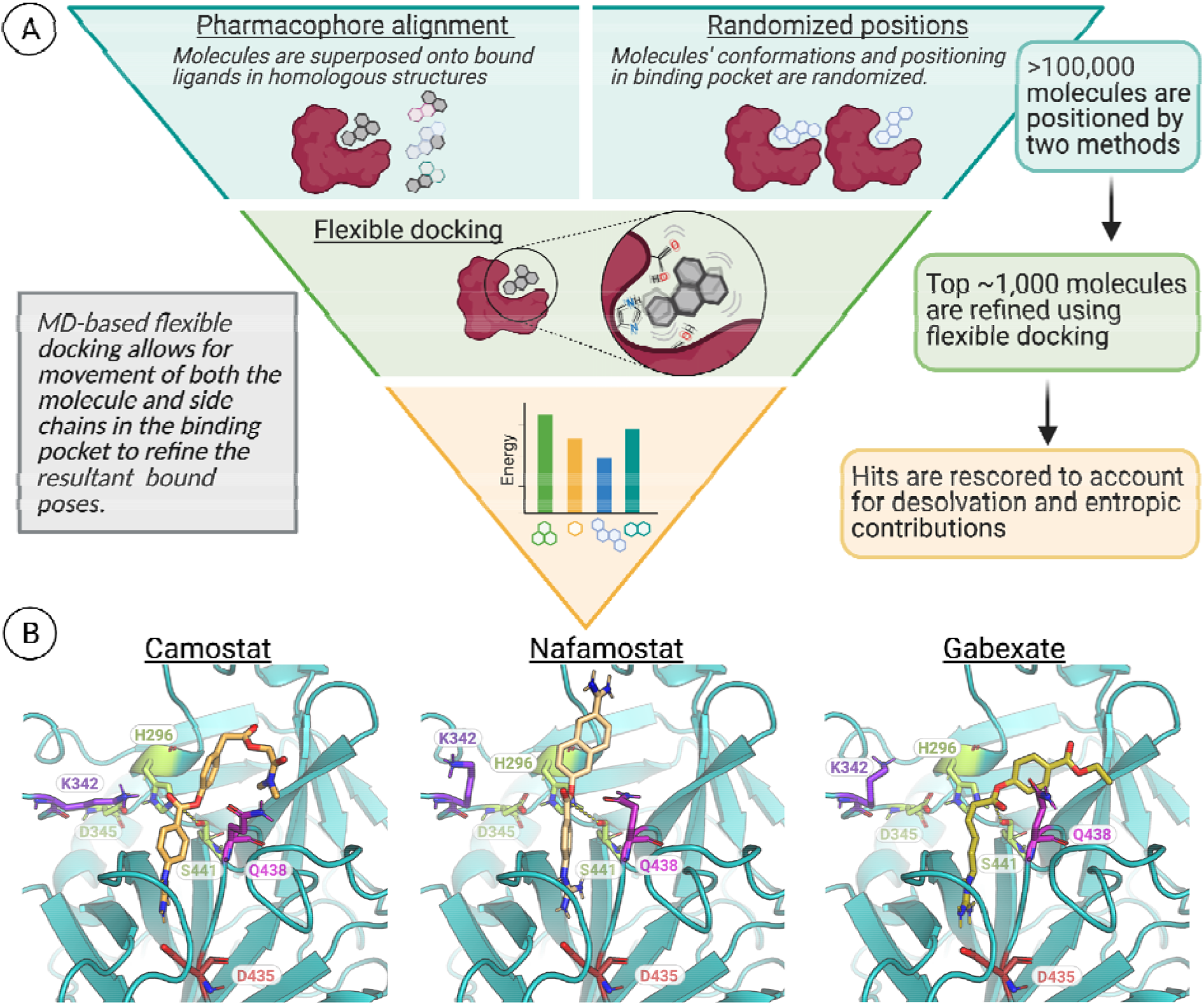
(A) The virtual screening protocol involves multiple refinements, including using MD-based docking. (B) Docked poses of camostat, nafamostat, and gabexate in the TMPRSS2 active site with the proton transfer from Ser-441 to His-296 that occurs after substrate binding. Shown in red is D435, which resides at the bottom of the S1 binding pocket, and lime green corresponds to the catalytic triad (His-296, Asp-345, Ser-441). Dashed lines shown here indicate the distance between the catalytic serine oxygen and the carbonyl carbon of each inhibitor (camostat = 3.5 Å, nafamostat = 4.8 Å, gabexate = 3.5 Å). Figure created using BioRender.com.

From this initial stage, 4,307 Level 1 screening candidates were determined and subjected to GPU accelerated Flexible-CDOCKER methods that were recently developed as part of the CHARMM molecular modeling package cite.^45–48^ This approach utilizes flexible side chains for residues in or near the binding pocket while using a grid representation for the remaining receptor. Multiple copies of each set of side chains and initial ligand poses are created, which allows for parallel, multiple copy processing of multiple flexible ligands-flexible receptor trials simultaneously on GPUs. The flexible docking searching algorithm combines molecular dynamics (MD) based simulated annealing and a continuous genetic algorithm search protocol to enhance the sampling of differing receptor conformations.

We utilize a novel scoring methodology by performing conformational clustering of the flexible side chains and the ligand, which provides key contributions to the ligand scoring from the entropic variation of the side chains to accommodate various ligand poses. The ligands are rescored in the protein binding site a final time using an implicit solvent model that captures aspects of the desolvation costs not generally accessible in typical docking methods.^49^ The rescoring is accomplished by minimizing the docked poses from the flexible side chains and flexible ligand in the context of the rigid protein, while also considering the total energy of the solvated docked and undocked systems.

The virtual screen was successful on many fronts. MD-based flexible docking identified residues near the active site that are conformationally dynamic to accommodate different ligands. Residues Gln-438 and Lys-342 in particular show the greatest conformational change upon ligand binding, suggesting that they participate in stabilizing bound compounds. The known inhibitors camostat, nafamostat, and gabexate all ranked in the top 5 compounds, and all adopted poses that demonstrate how the catalytic serine residue positions itself to ultimately react with the inhibitors while positioning the guanidinium functionality to form a salt-bridge interaction with the active site Asp-435. As the mechanism of covalent inhibition involves His-296 deprotonating Ser-441, we performed a subsequent docking experiment using the three molecules with these charge changes (Figure 4B). For all three molecules, the deprotonated serine positions itself into a more reactive state to attack the carbonyl carbon. For instance, the distance between the serine oxygen and camostat carbonyl carbon decreased from 4.9 Å to 3.5 Å, and the distance for gabexate decreased from 5.9 Å to 3.3 Å. While nafamostat appears to be further away from the reactive carbon (4.4 Å to 4.8 Å), the molecule flips so that the carbonyl is positioned for reactivity.

### Identification of noncovalent inhibitors

Several clinically approved drugs emerged as top ranked compounds in the virtual screen, which we selected to test in our *in vitro* assay. Like the covalent inhibitors, pentamidine, propamidine, and debrisoquine all contain a guanidinium moiety and docked into the active site of TMPRSS2 with the positive charge pointing towards Asp-435 (Figure 5B). Biochemically, we found that all three molecules did in fact inhibit TMPRSS2 activity with varied potencies, with debrisoquine being the least potent (Figure 5A). The docked poses of pentamidine and propamidine show both compounds are positioned to block the active site residues, whereas debrisoquine does not fully span the catalytic triad, which likely correlates to the differences in potency (Figure 5B). Pentamidine and propamidine are of similar size to camostat and nafamostat, typical of small molecule inhibitors (>350 MW), while debrisoquine is quite small, at 175.2 MW, classifying it as a fragment rather than a small molecule. However, debrisoquine has the greatest ligand efficiency (LE) at 0.42 compared to pentamidine and propamidine, which are 0.33 and 0.31 respectively. Thus, for its small size debrisoquine binds well to TMPRSS2. Furthermore, a LE = 0.42 is suggestive of an excellent starting point for fragment expansion efforts. It is worth noting that each of these molecules also inhibit trypsin activity (Figure S7). However, the fragment debrisoquine has modest selectivity for TMPRSS2 over trypsin (~3 fold). As observed with the covalent inhibitors (camostat, nafamostat, and gabexate; Figure 4B), the docking studies with the noncovalent compounds showed significant structural rearrangements for the two non-catalytic residues, Lys-342 and Gln-438 (Figure 5B). While Gln-438 is conserved across the serine proteases tested here as well as those used to construct the homology model (i.e. TMPRSS2, trypsin, hepsin; lysine residue in human plasma kallikrein), Lys-342 only appears in TMPRSS2. In fact, the entire loop where this residue resides greatly differs in length and conformation among the four proteases. Thus, derivatization of a fragment like debrisoquine towards Lys-342 may confer additional selectivity across similar serine proteases.

**Figure 5.**
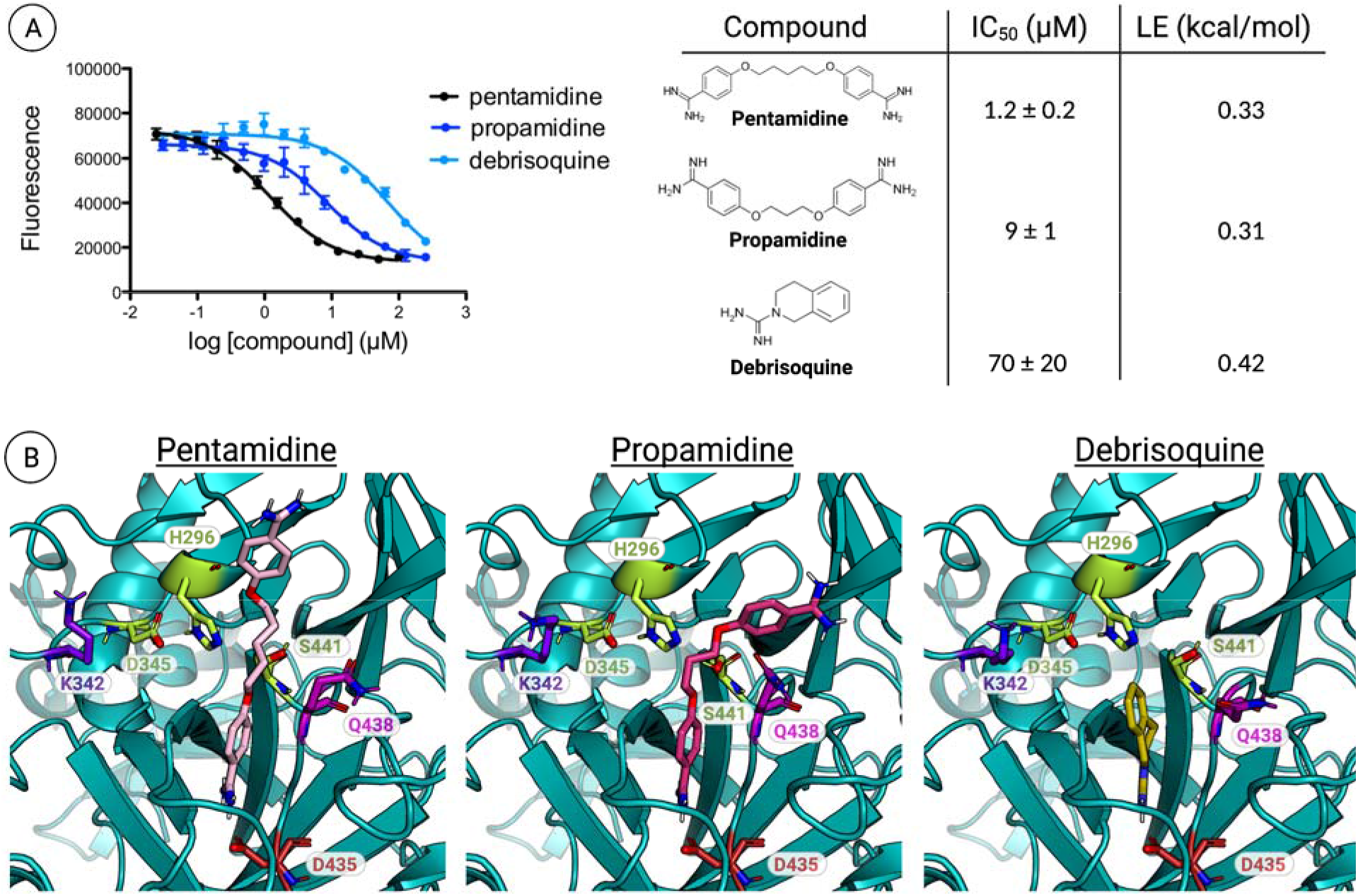
(A) Computational hits inhibit TMPRSS2 in biochemical activity assay. Left: Raw inhibition values used to obtain IC_50_ curves. Data is the average of duplicate experiments conducted in technical triplicate. Right: Table showing structures of hits, calculated IC_50_ values, and ligand efficiencies (LE). (B) Docking results for the three drugs identified as hits both in the virtual screen and the *in vitro* assay. All three molecules fit into the active site (Asp-435 at the bottom of the pocket shown in red). Pentamidine and propamidine obstruct access to the catalytic triad (shown in lime green), whereas the fragment debrisoquine only partially reaches those residues. Figure created using BioRender.com.

## Conclusion

With new strains of SARS-CoV-2 emerging, and the significance of TMPRSS2 in viral entry for multiple coronaviruses, it is pivotal that we uncover novel strategies to inhibit TMPRSS2 protease activity.^50,51^ However, TMPRSS2 provides obstacles in multiple areas of the discovery pipeline. For instance, the lack of an experimentally solved structure makes docking studies a challenge, relying instead on the use of a homology model developed based on other serine protease domains. Biochemically, the protease domain of TMPRSS2 has proven difficult to purify and refold recombinantly in *E. coli*, which has stalled many high-throughput screening efforts. In the present study, we demonstrate that combined computational and experimental techniques can be used to identify new TMPRSS2 protease inhibitors. Having identified promising scaffolds with high ligand efficiencies, future work will be dedicated towards improving potency and selectivity of these inhibitors. While we have developed an effective expression and purification protocol for the TMPRSS2 peptidase domain, it remains a challenge to obtain high yields of active protein. Thus, the combined virtual and biochemical screening approach presented here is attractive because it enables an initial triage through large compound libraries before testing a smaller number of molecules, more likely to be functionally relevant, in an assay. Current efforts are directed toward testing more hits prioritized from the computational screen for biochemical activity.

## Methods

### Mass spectrometric analysis of trypsin

Trypsin (Sigma, T9201), 10 μM, was incubated with compound, 10 μM, for 30 minutes at room temperature in assay buffer (50 mM Tris, 500 mM NaCl, 0.001% NP-40, pH 7.5) before being subjected to analysis by mass spectrometry using an Agilent QToF LC-MS equipped with a Poroshell 300SB C8 reverse phase column. A 5-100% gradient of acetonitrile with 0.1% formic acid in water to 0.1% formic acid over five minutes was used. Raw data was deconvoluted (intact protein of 20,000-25,000 Da) using BioConfirm software with background subtraction.

### Vector design

Plasmids were constructed by Twist Bioscience by inserting the gene for protease domain of TMPRSS2, specifically amino acids 247-492, into the pET28a+ vector using the NdeI_XhoI restriction enzyme cut-sites.

### Protein expression

The pET28a+ plasmids containing the TMPRSS2 genes were transformed into BL21(DE3) and plated on LB agar with kanamycin. The bacteria were grown in small 5 mL LB (+ kanamycin) cultures overnight at 37 °C. The 5 mL starters were used to inoculate 1 L LB (+ kanamycin) cultures, which were grown to OD = 0.8 at 37 °C with shaking at 250 rpm. Expression was induced using 1 mM Isopropyl β-d-1-thiogalactopyranoside (IPTG), which we let grow for 5 hours. The cells were then spun down at 9,500 x g for 15 min. The pellets were collected, flash frozen and stored at −80 °C.

### Chemical lysis and denaturing

Before lysing, the cell pellet was first fully thawed until it reached room temperature. Chemical lysis was performed by resuspending the pellet using B-PER reagent (Fisher, PI78243) with lysozyme (Fisher, 90082) and DNase I (Fisher, 90083) following manufacturer’s protocols. The cells were then spun at 15,000 x g for 5 minutes, and the cell lysate was collected and saved. The insoluble portion, which contained inclusion bodies of TMPRSS2, was resuspended / washed using lysis buffer containing detergent (50 mM Tris HCl, 0.9% NaCl, 1% Triton X-100, pH 7.5) and then spun at 15,000 x g for 5 minutes. After removing the supernatant, the pellet was washed once more with a lysis buffer that did not contain detergent (50 mM Tris HCl, 0.9% NaCl, pH 7.5) and then spun at 15,000 x g for 5 minutes.

The pellet was resuspended and the inclusion bodies were denatured by adding 20 mL denaturing buffer (8 M urea, 10 mM Tris, 100 mM sodium phosphate, pH 8.0) plus reducing agent (1:1000 BME). Denaturing occurred at room temperature with rotation for at least 30 minutes. The concentration of protein was determined via nanodrop, and additional denaturing buffer was added to reduce the concentration to below 1mg/mL. Denaturing occurred at room temperature on a rotator for at least 30 minutes before being spun down and decanted (20,000 x g, 15 minutes).

### Batch binding

Ni-NTA agarose (Qiagen, 30210) was prepared by washing 3 times with binding buffer (8 M urea, 10 mM Tris, 100 mM sodium phosphate, pH 8.0). Denatured protein was added to Ni-NTA resin (750 μL) and incubated at 4 °C on a rotator for 1.5 hours. Resin was pelleted by centrifugation at 2500 x g and flowthrough was removed. Resin was washed 3 times with wash buffer (8 M urea, 10 mM Tris, 100 mM sodium phosphate, 20 mM imidazole, pH 6.5), followed by addition of elution buffer (8 M urea, 10 mM Tris, 100 mM sodium phosphate, 500 mM Imidazole, pH 6.5). Eluting was performed on a rotator at 4 °C for 30 minutes. The resin was again pelleted by centrifugation at 2500 x g, and the sample was collected.

### Refolding

The denatured sample was diluted 1:100 into refolding buffer (50 mM Tris, 0.5 M arginine, 20 mM CaCl_2_, 1 mM EDTA, 100 mM NaCl, 0.01% NP-40, 0.05 mM GSSG, 0.5 mM GSH, pH 7.5) at room temperature using a syringe pump (flow rate 1 mL/min) while allowing the solution to gently stir. The refolding protein was left at 4 °C for 3 days with gentle stirring.

The sample was concentrated 10-fold using Amicon Stirred Cells with 10 kDa Ultrafugation disks (Millipore, UFC801024). Once concentrated, the sample was dialyzed overnight into assay buffer (50 mM Tris, 500 mM NaCl, 0.001% NP-40, pH 7.5) at 4 °C.

### Protein gel and silver staining

LDS loading dye was added to protein samples and samples were boiled for 5 minutes at 95 °C. 10uL of each sample was loaded onto a 4-20% mini-PROTEAN TGX gel (BioRad, 4561096) and run at 180V for 45 minutes. Total protein was visualized using a Pierce Silver Staining Kit (Thermo, PI24612) following manufacturer’s protocols.

### Western Blot

After running gel as described above, protein was transferred to PVDF membrane using a BioRaD TransferBox Turbo following the standard protocols. Membrane was blocked for 1 hour at room temperature using Super Block (Thermo Scientific, 37515). TMPRSS2 antibody (Novus biologicals, NBP1-20984) was added to membrane (1:1000 dilution in Super Block) and incubated overnight at 4C with gentle shaking. After removal of primary antibody and three washes with TBST, HRP conjugated secondary antibody (abcam, ab6741, 1:20,000 in Super Block) was added to membrane and incubated at RT for 1hr with shaking. After removal of secondary antibody with three washes with TBST, HRP substrate (Thermo Scientific, 34095) was added and after 1 minute Western blot was visualized using Chemiluminescence on an Azure Biosystems c600 imager.

### Kinetic assays

Assays were conducted on a Molecular Devices Spectramax Spectrophotometer using 96-well plates (Fisher, 12-565-501). Protein was first plated, followed by addition of substrate, Boc-QAR-AMC (Bachem, 4017019.0005) at concentrations to give the indicated final concentration in a 100 μL volume. After addition of substrate, fluorescence was immediately read (Ex: 380, Em: 460nm), taking measurements every 30 seconds for 20 minutes. Active protein was quantified by titrating in the known active site protease inhibitor FPR-CMK (Haematologic Technologies). To determine the K_M_ the initial fluorescence data, at less than 10% substrate conversion, was fit to a linear equation and the slope was determined, V_0_. V_0_ was plotted vs substrate concentration and the data was fit to the Michaelis Menten equation using GraphPad Prism.

### Fluorescence endpoint assays for IC_50_ determination

Assays were conducted in 384 well black plates (Costar, 4514) using an Envision plate reader, ex. filter 350 nm and em. filter 450 nm. The compounds were first plated (10 μL, at various concentrations) followed by addition of TMPRSS2 protein (8 μL, 0.5 nM final concentration). After 30 minute incubation (unless otherwise specified) at room temperature, substrate (2 μL, 2.5 μM final concentration) was added. At 30 minutes, corresponding to less than 20% substrate cleavage as measured by comparing fluorescence of the negative control to free AMC (Millipore, 257370), fluorescence was read. Wells containing no TMPRSS2 protein (substrate only) served as positive controls. Wells containing no inhibitors (TMPRSS2 and substrate only) served as negative controls. Fluorescence readout was plotted against the log of inhibitor concentration and fit to log(inhibitor) vs response - variable slope equation in GraphPad Prism. Fluorescence endpoint assays with trypsin were conducted utilizing 1 nM protein and 5 μM substrate.

### General flexible docking setup

The homology model of TMPRSS2 was generated using SWISS-Model based on the serine protease Hepsin (PDB 5CE1), which has 34% similarity and 70% coverage of the TMPRSS2 sequence. Included in the Hepsin structure is a 100 nM inhibitor, 2-[6-(1-hydroxycyclohexyl)pyridin-2-yl]-1H-indole-5-carboximidamide, which is bound in the active site. The inhibitor is utilized as one of the pharmacophore targets in our *fastdock* protocol.

The *fastdock* protocol is a python-based workflow that integrates the *align-it* software to search across our curated library of compounds for 3D pharmacophore matches to an inhibitor from a solved structure.^52,53^ The *fastdock* ligand templates are taken from the Hepsin structure used in the initial generation of the model (PDB 5CE1) as well as from a plasma kallikrein structure with the 1 nM inhibitor N-[(6-amino-2,4-dimethylpyridin-3-yl)methyl]-1-({4-[(1H-pyrazol-1-yl)methyl]phenyl}methyl)-1H-pyrazole-4-carboxamide bound (PDB 6O1G; 43% sequence similarity and 51% sequence coverage). Scoring of the pharmacophore matches is based on a volumetric Tanimoto value of the target ligand pharmacophore map and the reference ligand map. Based on this initial selection of potential ligands for exploration, we harvested 1-10% of the top hits.

The MMTSB tool set was used to cluster binding poses and prepare pdb files.^54^ Open Babel was used to generate ligand random conformations.^55^ MOE was used to predict the correct protonation state for the ligands at pH 7.4.^56^ ParamChem was used to prepare the ligand topology and parameter files with the CGenFF force field.^57–59^ Clustering used the tool cluster.pl with a 1 Å cutoff radius for the K-means clustering. The CHARMM C36 force fields were used and docking was performed in CHARMM with the CHARMM/OpenMM parallel simulated annealing feature.^48,60^

### Flexible docking setup

Flexible CDOCKER with a hybrid searching algorithm combining molecular dynamics (MD) based simulated annealing and continuous genetic algorithm was used to dock and rank the top hits.^45^ Flexible CDOCKER uses a physics-based scoring function and allows both ligand and protein side chains to explore their conformational space simultaneously. The following amino acid side chains are considered flexible: His-296, Tyr-337, Lys-342, Asp-435, Ser-436, Gln-438, Ser-441, Thr-459, Trp-461 and Cys-465.

Each docking measurement represents 500 genes (docking trials). The coordinates of the ligand-protein flexible side chains are used to assemble a gene (potential docking pose). Each ligand in the dataset is first aligned to the pharmacophore model with *align-it*. In the initial generation, half of the genes have the ligand starting with the aligned position. The rest of the genes are constructed by generating a random conformation of the ligand with Open Babel and centering at the binding pocket. A random translation (within a volume with a 2 Å edge length) and rotation (maximum 360○) are performed on ligands in each gene. An energy cutoff is applied to avoid potential collision between ligand atoms and protein atoms due to the random translation and rotation. The protein flexible side chains are initialized with the coordinates from the input homology model. Then these genes are optimized by an MD based simulated annealing algorithm.

Detailed values for softness parameter *E_max_* used in flexible receptor docking are summarized in Table 1.

**Table 1:**
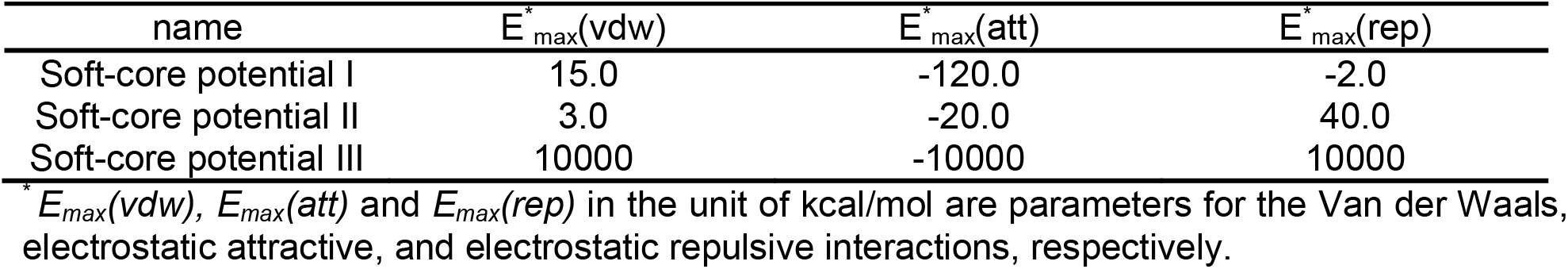
Soft-core potentials used in flexible receptor docking

| name | $E_{max}^{*}(vdw)$ | $E_{max}^{*}(att)$ | $E_{max}^{*}(rep)$ |
| --- | --- | --- | --- |
| Soft-core potential I | 15.0 | -120.0 | -2.0 |
| Soft-core potential II | 3.0 | -20.0 | 40.0 |
| Soft-core potential III | 10000 | -10000 | 10000 |

\* $E_{max}(vdw)$ , $E_{max}(att)$ and $E_{max}(rep)$ in the unit of kcal/mol are parameters for the Van der Waals, electrostatic attractive, and electrostatic repulsive interactions, respectively.

The docking poses (optimized genes) are then K-means clustered based on ligand heavy atom RMSD with a radius cutoff of 1 Å. We then select the best individuals (minimum energy pose) from the top 10 largest clusters to construct the second generation. In our previous study, we show that using two generations is adequate and the average computer time for each docking measurement is around 30~45 mins. After the second generation, the docking poses are clustered and the best individuals from the top 15 largest clusters are saved. These docking poses are then rescored using the FACTS implicit solvent model.^49^

## Acknowledgements

We want to acknowledge the valuable scientific input from our entire TMPRSS2 team, including Dr. Corey Stephenson, Kevon Stanford, Tessa Epstein, Dr. Stephen Joy, Furyal Ahmed, Dr. Allison Narayan, and Dr. Madison Sowden. Research reported in this publication was supported by the Center for Chemical Genomics (CCG) at the University of Michigan Life Sciences Institute. Financial support for this work was provided by the Klatskin-Sutker Discovery Fund award (to A.K.M.) and the National Institutes of Health Grants GM103695 (to C.L.B) and GM130587 (to C.L.B).

## Table of contents graphic

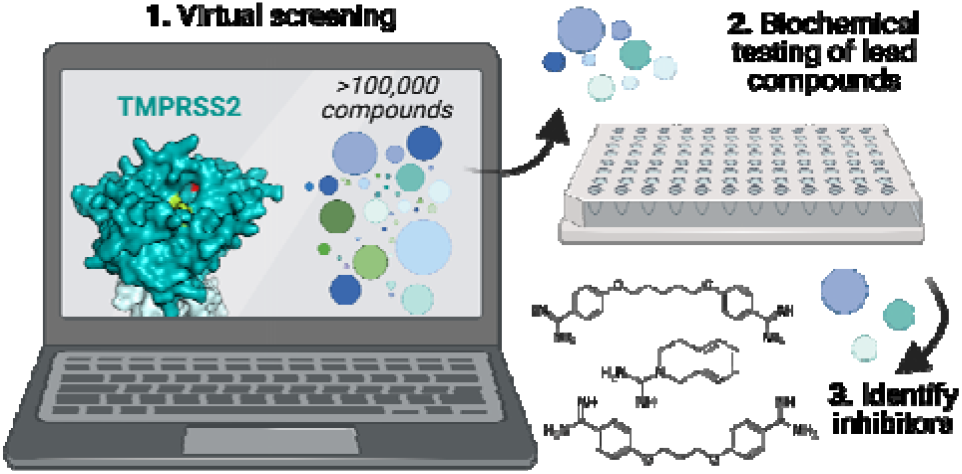

