## Supplemental materials for "TMPRSS2 inhibitor discovery facilitated through an *in silico* and biochemical screening platform"

Table of Contents:

Figure S1. Western blot for 6xHis tag at various stages of TMPRSS2 purification

Figure S2. Concentration of TMPRSS2 active protein calculated using FPR-cmk

Figure S3. $K_M$ calculation for Boc-QAR-AMC with Trypsin

Figure S4. Inhibition curves of camostat, nafamostat, FOY 251, and gabexate obtained using TMPRSS2 biochemical assay

Figure S5. LC-MS data of showing adduct formation of nafamostat and FOY 251 with trypsin

Figure S6. LC-MS data showing covalent adduct formation of gabexate with trypsin

Figure S7. Inhibition of Trypsin by pentamidine, propamidine, and debrisoquine

Table S1. Ranking of compounds from computational screening

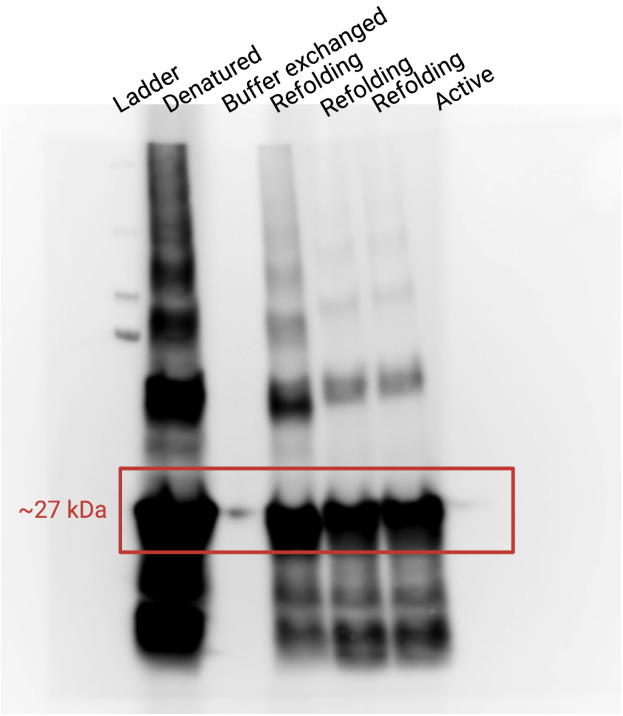


**Figure S1.** Western blot for 6xHis tag at various stages of TMPRSS2 purification. Significant signal is seen after denaturation and at various time points during refolding. A small amount of His-tagged protein is observed immediately after exchange of concentrated refolding solution into activation buffer, but ultimately no signal is observed, indicating autocleavage and removal of 6xHis tag, in the final active product.

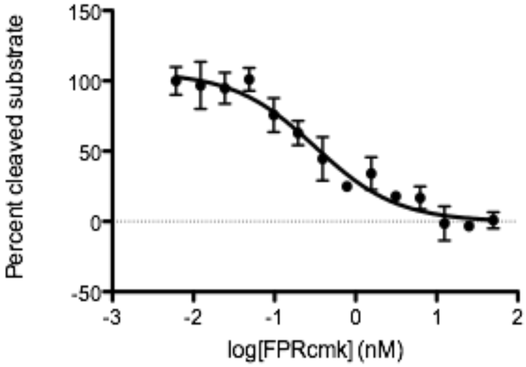


**Figure S2**. Inhibition curve of TMPRSS2 activity obtained with increasing [FPR-cmk]. The $IC_{50}$ obtained here is 0.32 ± 0.09 nM, allowing us to approximate 100% inhibition at about 0.64 nM. Thus, assuming 1:1 complex formation of protein and inhibitor, we can determine the protein concentration of this sample to be about 0.64 nM. Data points represent the average of technical triplicate data points with error bars indicating the standard deviation.

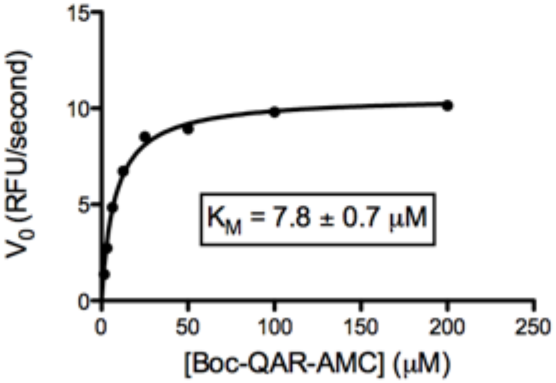


**Figure S3.** Plot of the initial rate vs substrate concentration for trypsin with Boc-QAR-AMC. Fitting to the Michaelis-Menten equation gives the $K_M$.

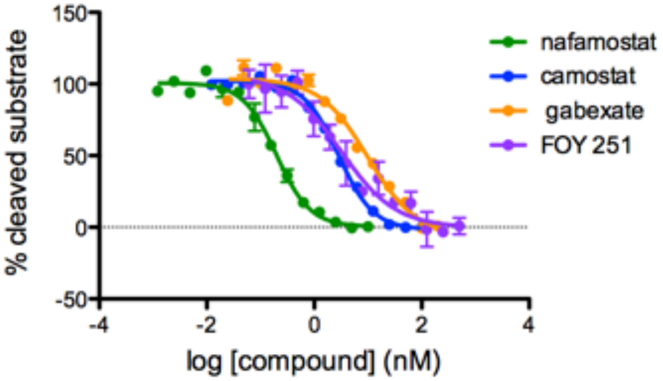


**Figure S4.** Inhibition of TMPRSS2 by compounds camostat, nafamostat, FOY 251, and gabexate, with 30 minutes preincubation of protein and inhibitor. Data points represent the average of technical triplicate with error bars indicating the standard deviation.

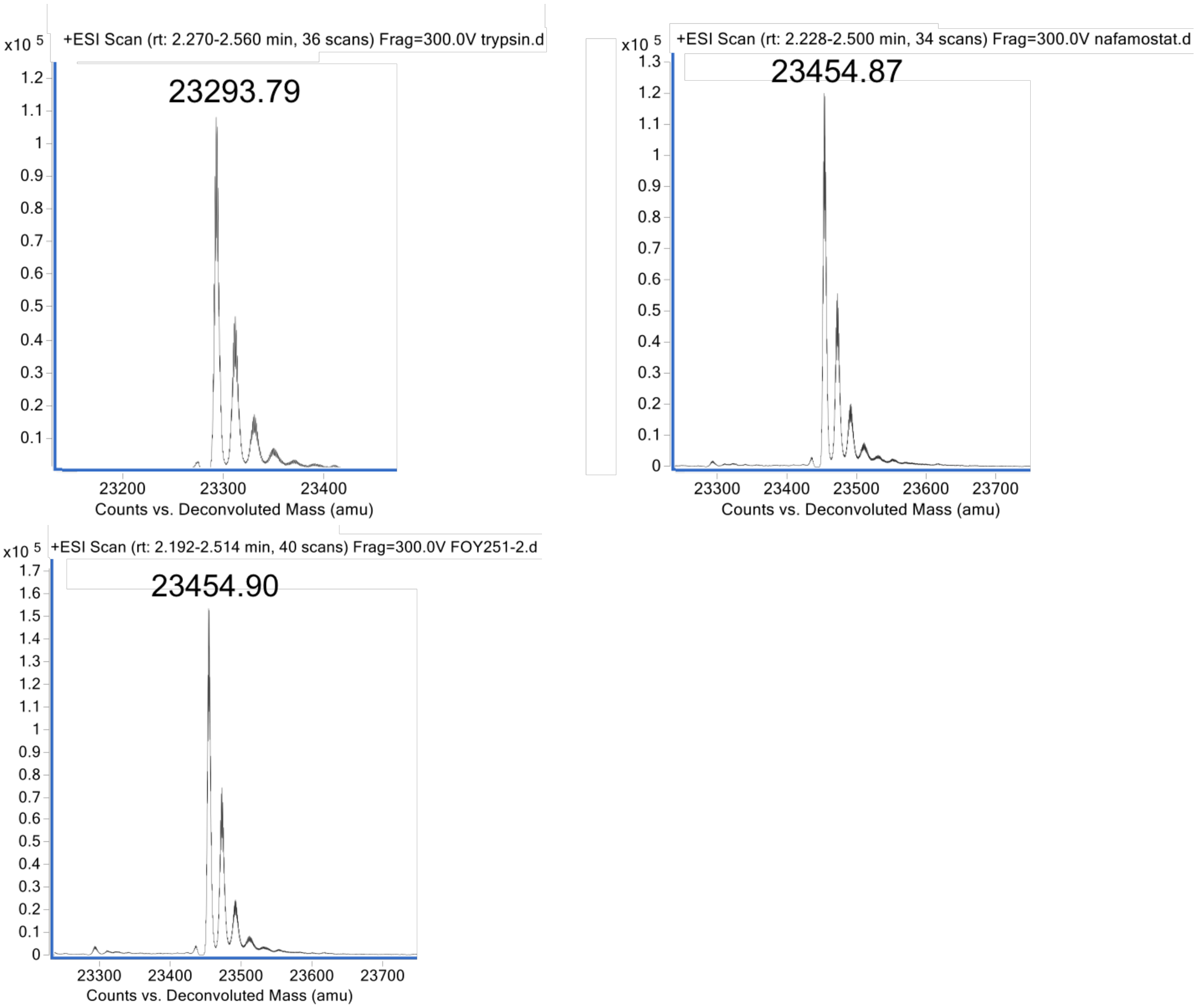


**Figure S5.** Deconvoluted LC-MS data of trypsin (top left), incubated with equimolar nafamostat (top right), or FOY 251 (bottom). After 30 minutes incubation, a mass increase of about 161 Da is observed, indicating covalent adduct formation.

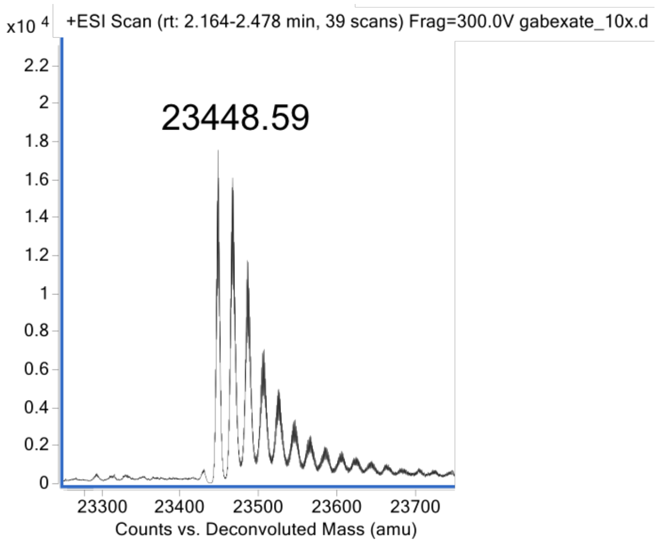


**Figure S6.** Deconvoluted LC-MS data of trypsin incubated with 10X gabexate. After 30 minutes incubation, a mass increase of about 155 Da is observed, indicating covalent adduct formation. No mass change was observed after 30 minutes incubation with equimolar inhibitor.

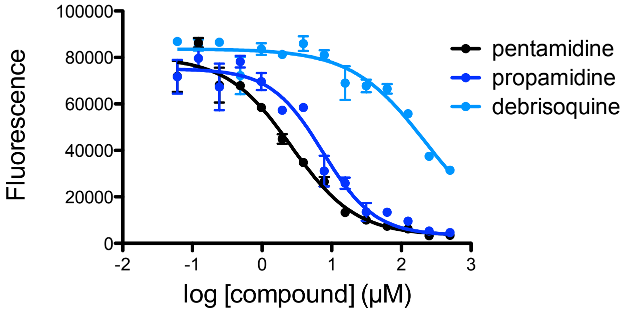


| | $IC_{50}$ (μM) |
|---|---|
| debrisoquine | 220 ± 30 |
| pentamidine | 2.6 ± 0.4 |
| propamidine | 7 ± 1 |

**Figure S7.** Inhibition of Trypsin by computational hit compounds pentamidine, propamidine, and debrisoquine with 30 minutes preincubation of protein and inhibitor. Data points represent the average of technical triplicate with error bars indicating the standard deviation (left). Calculated $IC_{50}$ values (right).

**Table S1**. Top ranked molecules from the TMPRSS2 virtual screen. Libraries screened included the ZINC database,[1] SWEETLEAD,[2] as well as compounds from the Center for Chemical Genomics (CCG) at the University of Michigan.

| Compound ID | Compound name | Rank |
|---|---|---|
| ZINC000003594299 | Debrisoquine | 1 |
| Gabexate | Gabexate | 2 |
| ZINC000003871842 | Camostat | 3 |
| ZINC000473158220 | | 4 |
| ZINC000003874467 | Nafamostat | 5 |
| ZINC000015555321 | | 6 |
| ZINC000095558579 | | 7 |
| ZINC000001532560 | | 8 |
| ZINC000006092735 | | 9 |
| ZINC000001665564 | Propamadine | 10 |
| CCG-335777 | | 11 |
| CCG-335408 | | 12 |
| ZINC000004642990 | | 13 |
| ZINC000001530775 | Pentamidine | 14 |
| CCG-294352 | | 15 |
| SW03040 | | 16 |
| CCG-302966 | | 17 |
| CCG-358814 | | 18 |
| ZINC000029130869 | | 19 |
| CCG-320104 | | 20 |
| CCG-358814 | | 21 |
| ZINC000008101126 | Guanidine | 22 |
| ZINC000000599985 | | 23 |
| CCG-335801 | | 24 |
| CCG-355904 | | 25 |
